# Behavioral evidence challenges species-specific ocular morphology as a primary constraint on human gaze-following

**DOI:** 10.64898/2026.05.11.705002

**Authors:** Masih Shafiei, Yasmin Arnous, Nick Taubert, Martin Giese, Peter Thier

**Affiliations:** Cognitive Neurology Lab, Hertie Institute for Clinical Brain Research, Eberhard Karls University Tübingen, Hoppe-Seyler-Str. 27, 72076 Tübingen, Germany; Graduate Training Centre of Neuroscience, International Max Planck Research School, Eberhard Karls University Tübingen, Hoppe-Seyler-Str. 27, 72076 Tübingen, Germany; Faculty of Medicine and Faculty of Mathematics and Natural Sciences, University of Cologne, 50931 Cologne, Germany; Section Computational Sensomotorics, Department N3, Hertie-Institute for Clinical Brain; Werner Reichardt Centre for Integrative Neuroscience, Eberhard Karls University Tübingen, Otfried-Müller-Straße 25, 72076 Tübingen

**Keywords:** Gaze-following, social attention, visual spatial attention, eye morphology

## Abstract

Previous research suggests that humans are extremely sensitive to object-directed eye gaze, which effectively guides their attention toward objects of shared interest. This contrasts with non-human primates, who typically require much more salient eye-gaze cues to achieve comparable attentional orienting. However, it remains unclear whether cross-species differences in ocular morphology account for this performance gap. To address this question, we examined humans’ covert shifts of spatial attention in response to eye-gaze cues provided by either realistic human or rhesus monkey head avatars. Target detection was reliably enhanced on gaze-congruent compared to gaze-incongruent trials, with comparable gaze-cueing effects for both avatar types, despite the fact that monkey eyes lack many of the conspicuous features characteristic of human eyes. Hence, eye morphology alone does not substantially modulate gaze-driven attentional orienting in humans, whereas humans’ reliable use of monkey eye-gaze cues highlights a clear species difference in perceptual sensitivity to eye gaze signals.

**Significance Statement:** Eye-gaze–mediated attentional orienting is a conserved ability across primates, yet sensitivity to subtle eye-gaze cues may differ between species. Here, we provide empirical evidence that humans exhibit a quantitatively greater capacity than non-human primates to follow subtle eye-gaze cues. Importantly, we showed that this difference cannot be attributed to species-specific ocular morphology as human participants showed robust and comparable reflexive attentional orienting to both human and rhesus monkey eye-gaze cues. This is striking given the pronounced differences in ocular morphology and coloration/contrast between the two species. These findings suggest that cross-species diversity in extracting spatial information from eye-gaze cues likely reflects differences in perceptual sensitivity rather than bottom-up constraints imposed by species-specific ocular morphology.

## Introduction

Human communication relies heavily on language, yet a substantial portion of socially relevant information is conveyed through nonverbal signals that engage visual and motor systems. Among these signals, gaze direction—defined by the orientation of the eyes and influenced by the orientation of the face or body—plays a particularly prominent role by indicating where another individual’s attention is directed. The ability to co-orient oneself with another’s gaze direction, known as gaze-following (1), enables observers to detect behaviorally relevant aspects of the environment, such as threats or resources, that might otherwise go unnoticed. Beyond these immediate perceptual benefits, gaze-following also supports higher-level social cognition by enabling individuals to infer what others see as well as their goals, intentions, and mental states shaped by their objects of interest—a route to the development of a *theory of* (the other’s) *mind* (2, 3).

Examining how gaze-following operates across species is critical for determining which aspects of this ability reflect shared, evolutionarily conserved mechanisms and which may depend on species-specific adaptations. Non-human primates (NHPs) are of particular relevance to this question given their close evolutionary relationship with humans and the homology in visuo-oculomotor system organization (4, 5) which could in principle support perceptual sensitivity to fine visual details comparable to that of humans. Until recently, the dominant view held that humans possess a categorically superior gaze-following capacity (6), often attributed to the exceptional conspicuity of their eyes. According to this view, eye-gaze following was considered uniquely human and rooted in distinctive ocular morphology (7, 8). However, more recent comparative morphological analyses have undermined this latter claim, showing that features putatively allowing the perception of gaze direction are also present in several non-human primate species and, moreover, that substantial intra-species variability exists in scleral pigmentation and eye contrast within humans (see Perea-García et al.(9) for a review). While these findings substantially weaken a strictly human-specific morphological account, it remains unclear whether these anatomical differences translate into variation in gaze-following behavior.

To move beyond morphological description and directly assess functional relevance, we designed and conducted the present study. We asked whether gaze-following in humans is modulated by species-typical ocular morphology. To this end, we employed a gaze-cueing paradigm in which non-predictive gaze cues, presented by either ultra-realistic human or monkey avatars, preceded a brief, near-threshold luminance change at one of two peripheral target locations that participants were required to detect. Crucially, gaze cue were non-predictive within the task structure and participants were explicitly instructed that gaze did not signal target location, following gaze conferred no net strategic advantage for target selection. This design therefore allowed us to isolate and assess the spontaneous influence of gaze cues, independent of their instrumental value. Eye-gaze shifts demonstrated by the human and monkey avatars were matched in their trajectory and dynamics, thereby isolating the impact of species-typical ocular morphology to gaze-following. As expected, we observed significantly higher detection accuracy for targets appearing at gaze-congruent locations relative to incongruent locations, consistent with a robust gaze-cueing effect. Importantly, the magnitude, direction, and temporal dynamics of this effect were comparable for human and monkey avatars. These findings indicate that species-typical differences in ocular morphology do not substantially influence gaze cueing in humans under the present conditions. Instead, they suggest that humans’ robust ability to utilize eye-gaze cues reflects a species-specific advantage in perceptual sensitivity to gaze signals.

## Results

Hit probability (hit vs. miss) was analyzed using a generalized linear mixed-effects model with a binomial error distribution and logit link. Fixed effects included avatar type (human, monkey), congruency (congruent, incongruent), SOA (50, 100, 200, 300, 400 ms), and their interactions. Random intercepts and random slopes for congruency were included for each subject to account for the variability in baseline performance as well as the slope of the congruent-incongruent association across participants (figure 1a and table S1).

**Figure 1.**
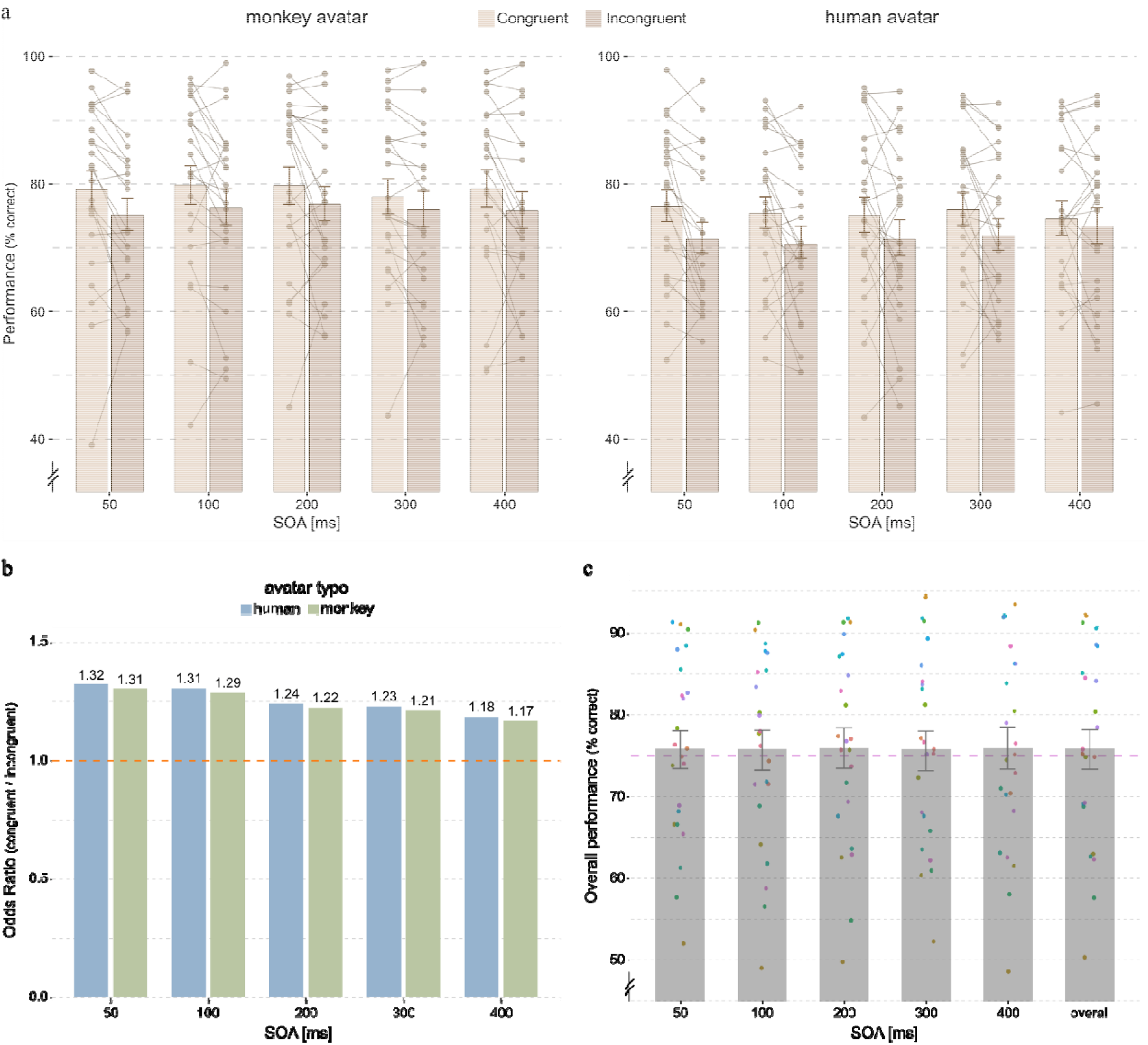
**(a)** Mean performance (% correct) as a function of stimulus onset asynchrony (SOA) for the monkey avatar (left) and human avatar (right), shown separately for congruent and incongruent gaze conditions. Bars represent group means ± SEM, and thin lines connecting pairs of datapoints depict individual subjects’ paired performance across congruency conditions. **(b)** Odds ratios (ORs) for congruent relative to incongruent trials derived from the mixed-effects model, shown as a function of SOA for monkey and human avatars. Values above bars indicate ORs; the dashed horizontal line at OR = 1 denotes no gaze-cuing effect. **(c)** Overall performance collapsed across congruency conditions and avatar variants (± SEM). Mean accuracy across SOAs was close to the targeted perceptual threshold of 75% (dashed line), indicating that task difficulty was successfully calibrated to maintain comparable performance levels across conditions. Color-coded individual subject data are overlaid to illustrate between-subject variability and allow tracking of individual average performance across SOAs.

We found significant main effects of avatar type (human vs. monkey) and congruence (congruent vs. incongruent) on hit probability (see table S1 for details). Participants showed overall lower odds of a correct response when viewing human avatars than monkey avatars (β = −0.232, SE = 0.032, z = −7.18, p < .001). This indicates a baseline performance difference due to type of avatar. The analysis demonstrated a higher hit probability on congruent compared to incongruent trials (β = 0.267, SE = 0.083, z = 3.22, p = .001), corresponding to increased odds of a correct response under congruent conditions. We refer to this effect as gaze-cuing effect. In contrast, SOA, interaction between SOA and congruency and interaction between avatar type and congruency had no significant effect on hit probability (all ps > .14). This indicates that overall performance did not systematically change across SOA levels when collapsing across congruency and species. Moreover, it showed that the congruency effect did not reliably vary as a function of SOA or avatar type. Thus, although overall hit probability differed between avatar types, both types benefited similarly from congruent relative to incongruent trials.

Post-hoc comparisons showed that for the monkey avatar, congruent trials were associated with significantly higher odds of a correct response at SOAs of 50–300 ms (all adjusted ps ≤ .022; figure 1a left panel and figure 1b). The gaze cuing effect decreased gradually with increasing SOA, with odds ratios ranging from 1.31 at 50 ms to 1.21 at 300 ms. At SOA of 400 ms, the congruency effect was no longer statistically significant after correction (OR = 1.17, padj = .062). For human avatars, congruent trials likewise yielded significantly higher odds of a correct response across all SOAs (all adjusted ps ≤ .044; figure 1a right panel). Odds ratios ranged from 1.32 at 50 ms to 1.18 at 400 ms, indicating a clear but gradually attenuating congruency benefit with increasing SOA (figure 1b). This showed that the gaze cues, demonstrated by either human or monkey avatar, robustly evoked gaze-following behavior in human participants. Moreover, post-hoc comparisons of the congruency effect across species revealed no significant differences at any SOA. The ratio of ORs comparing monkey avatar to human avatar did not differ from unity (OR ratio = 1.01, z = 0.31, p = .76), indicating that the magnitude of the gaze-cuing effect was statistically indistinguishable between species across all SOAs. This result is consistent with the absence of a significant species × congruency interaction in the mixed-effects model. Importantly, this finding does not contradict the significant main effect of species observed in the model. The main effect of species reflects a difference in overall hit probability, with human avatars giving lower baseline accuracy than monkey avatars across conditions, independent of gaze congruency. In contrast, the post-hoc OR analysis specifically tests whether the relative benefit of congruent compared to incongruent gaze cues differs between species. Thus, while overall performance differed between human and monkey avatars, both avatar species yielded a comparable increase in hit probability in response to congruent gaze cues.

The random-effects structure indicated substantial between-subject variability in baseline accuracy, as reflected by the magnitude of variance in random intercepts (variance = 0.43, SD = 0.66). This indicates that subjects differed markedly in their overall probability of producing correct responses. In addition, there was meaningful between-subject variability in the congruency effect (magnitude and valence), as evidenced by non-negligible variance in the random slopes for congruency (variance = 0.075, SD = 0.27), indicating that individuals varied in how their performance was modulated by gaze congruency. The correlation between random intercepts and congruency slopes was very small (r = 0.01), suggesting that subjects’ baseline accuracy was largely independent of their congruency effect. In other words, participants who were generally more accurate were not systematically more (or less) influenced by congruent versus incongruent conditions.

We did not find any significant main effects of participant sex (male vs. female) or avatar presentation order (monkey-first vs. human-first) on hit probability, nor did these factors modulate the gaze-cuing effect (all ps > .05). Likewise, the sex of the human avatar (male vs. female) had no significant effect on baseline performance (p = .50) and did not reliably alter the magnitude of the gaze-cuing effect across SOAs, with the exception of a marginal interaction at the longest SOA (400 ms, p = .05). Across all models, a robust main effect of congruency was observed, confirming higher hit probability on congruent relative to incongruent trials, while the absence of significant interactions indicates that gaze-cuing effects were stable across participant sex, avatar sex, and presentation order.

To enable a direct quantitative comparison of gaze-cuing performance across species, human participants had been tested using the same gaze-cuing paradigm and experimental setup previously employed in rhesus monkeys (Shafiei et al., *in press*), including the identical small-amplitude eye-gaze cue demonstrated by a monkey avatar. These methodological considerations had been implemented to ensure full comparability between studies, allowing us to directly assess quantitative differences in gaze-following capabilities between humans and non-human primates. Figure 2 compares the magnitude of the gaze-cuing effects in human and monkey observers, expressed as odds ratios (ORs), as a function of stimulus onset asynchrony (SOA), gaze-cue variant, and tested observer species. ORs for three gaze-cue variants are included in this figure: a small-amplitude gaze cue demonstrated by a monkey avatar (used in both the current and the previous study), a small-amplitude gaze cue demonstrated by a human avatar to human observers (exclusively used in the present study), and a large-amplitude gaze cue demonstrated by a monkey avatar (employed in the previous study) to monkeys. The two bars shown in dark olive and dark lavender correspond to the identical small-amplitude monkey-avatar stimulus used in both the human and monkey experiments, enabling a direct cross-species comparison. As shown in Figure 2, rhesus monkeys did not exhibit a gaze-cuing effect in response to the small-amplitude gaze cue across SOAs, with ORs remaining close to unity (ranging between 1.00-1.13). In contrast, the same animals showed a robust gaze-cuing effect (ranging between 0.94-1.30) when presented with a large-amplitude gaze cue demonstrated by the monkey avatar (light lavender bars), suggesting that monkeys can follow eye-gaze when the cue is sufficiently salient. In humans, however, the small-amplitude gaze cue demonstrated by the monkey avatar reliably induced a gaze-cuing effect across all SOAs, with large ORs (ranging 1.17-1.31). Importantly, the magnitude of this effect did not differ meaningfully from that elicited by the small-amplitude gaze cue demonstrated by the human avatar, indicating comparable effectiveness of the two cue types in human observers.

**Figure 2.**
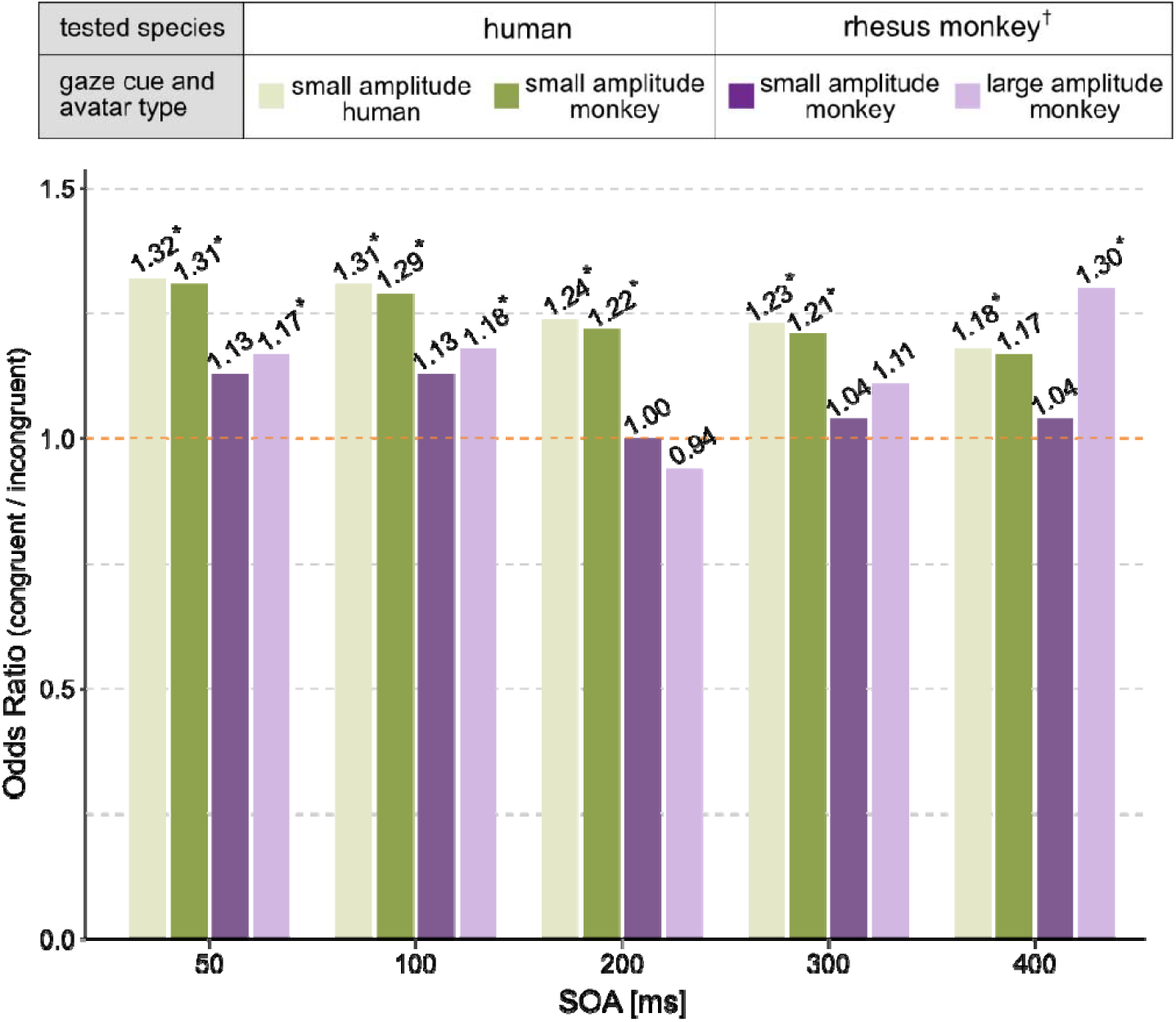
Cross-species comparison of gaze-cuing effect. ORs indexing the magnitude of the gaze-cuing effect are shown as a function of stimulus onset asynchrony (SOA), gaze-cue variant, and tested species (humans vs. rhesus monkeys). Three gaze-cue variants are included: a small-amplitude gaze cue demonstrated by a monkey avatar (dark olive and dark lavender), a small-amplitude gaze cue demonstrated by a human avatar (light olive), and a large-amplitude gaze cue demonstrated by a monkey avatar (light lavender). The light and dark olive bars correspond to gaze cues used in the present human experiment, whereas light and dark lavender bars represent conditions tested in our previous rhesus monkey study (Shafiei et al., in press). The dark olive (humans) and dark lavender (monkeys) bars correspond to the identical small-amplitude monkey-avatar stimulus, enabling direct cross-species comparison. ^†^Rhesus monkey ORs are replotted from Shafiei et al. (in press).

## Discussion

Previous research has shown that the ability to use conspecifics’ object-directed eye-gaze— independent of accompanying head movements—to orient one’s own attention toward an external object is a perceptual capacity shared by humans and non-human primates (see Shafiei et al., for a comprehensive review of NHP research, and Chac on-Candia et al., 2023 for a meta-analytic review of the human studies). Although these findings argue against a categorical, species-level dissociation, previous work has not been able to determine whether potentially relevant quantitative differences in this ability exist between humans and nonhuman primates (NHPs) and, if so, what mechanisms might underlie them. In the present study, we directly addressed this issue by comparing human gaze following induced covert shifts of visual spatial attention which were guided by either human or monkey eye gaze cues. Critically, we employed the same monkey gaze stimuli, behavioral paradigm, and even experimental setup we had previously used to study gaze-following in rhesus monkeys, thereby enabling a direct comparison of performance between monkey and human observers. More specifically, human participants viewed a realistic rhesus monkey head avatar – the same that had been used in the preceding study of rhesus monkeys - or, alternatively, a human head avatar, and were required to detect near-threshold luminance changes at one of two peripheral targets, with the demonstrator’s gaze directed toward the target location on 50% of trials, i.e. a non-predictive gaze cue. Target detection was significantly enhanced for gaze-congruent trials relative to incongruent trials, no matter if prompted by the human or the monkey avatar. In contrast, our prior work demonstrated that the identical monkey gaze stimulus was unable to induce gaze-following behavior in rhesus monkeys, who exhibited reliable eye gaze following only when prompted by a more salient cue characterized by increased eye-gaze deviation amplitude and reduced viewing distance. Taken together, these results support the existence of a clear quantitative species difference with humans exhibiting greater perceptual sensitivity to object-directed eye-gaze cues, allowing even low-quality visual input to elicit robust gaze-following effects. Moreover, the comparable efficacy of monkey and human gaze cues in eliciting human gaze following also challenges the widespread belief that differences in eye morphology explain previous observations of poorer eye gaze following in monkeys. This conclusion is allowed given the fact that the monkey and human avatar eyes reproduced the well-known differences in eye features. As shown in figure 2, the morphological distinctions are significant: the human palpebral fissure is characterized by a mediolateral elongation (almond-shaped), whereas the rhesus monkey aperture is markedly more circular. Furthermore, humans possess significantly greater scleral exposure across both primary and eccentric gaze positions. Regarding luminance-contrast relationships, brown-eyed humans exhibit a primary contrast boundary between the depigmented conjunctiva and the pigmented iris—rendering the pupil-iris transition less distinct—whereas rhesus monkeys (possessing hazel-brown irises and minimal visible sclera) present a dominant contrast between the relatively reflective iris and the dark pupil. Despite these profound configurational disparities, the magnitude, direction, and temporal dynamics of gaze-cueing effects exhibited by our human observers—as measured across stimulus-onset asynchronies (SOAs)—were nearly identical. These findings indicate that, for an average human observer, gaze cues derived from human and monkey eye morphologies are comparably salient and informative for guiding covert visual attention. This conclusion is in line with previous results from comparative studies of eye morphology across various non-human primate species and human populations (10–13), which have called into question the long-held belief that the seemingly poorer eye-following abilities of nonhuman primates might be a direct consequence of less conspicuous eyes as originally proposed by Kobayashi and Koshima (7, 8). In fact, collectively, more recent morphological work suggests that while clear differences in eye appearance exist, human eyes are not uniquely conspicuous (see Perea-García et al. (9) for a review).

We quantified gaze following using the cueing effect, defined as higher accuracy in detecting a luminance change when the target was preceded by a target-congruent gaze cue compared with a target-incongruent cue. This effect is interpreted as reflecting a covert shift of visual spatial attention guided by the avatar’s gaze. Covert orienting was ensured by task constraints: throughout gaze-cue presentation and until the go signal, participants were required to maintain fixation on a central fixation point, and any trial in which fixation was broken prior to the go signal was aborted and excluded from analysis. In incongruent trials, luminance changes were therefore more likely to be missed due to a misallocation of the attentional spotlight toward the distractor location cued by the gaze stimulus. Importantly, given the structure of the task, performance could not exceed 50% accuracy if participants relied exclusively on the gaze cue while ignoring target luminance; however, overall accuracy was approximately 75%, indicating that participants followed task instructions and reported luminance changes rather than simply following gaze. Moreover, gaze following was not explicitly reinforced, as auditory feedback signaling success was contingent solely on correct target detection, irrespective of cue congruency. Taken together, these considerations indicate that enhanced performance on gaze-congruent trials reflects spontaneous covert gaze-following tendencies, expressed despite the absence of explicit incentives or strategic benefits for the observer.

In the present experiment, the cue–target interval, or stimulus onset asynchrony (SOA), varied randomly across trials, with each SOA drawn with equal probability from a discrete set of five values (50, 100, 200, 300, and 500 ms). We observed that the gaze-mediated attentional effect emerged as early as 50 ms following cue onset. Such rapid modulation of performance is consistent with a reflexive orienting mechanism driven by exogenous properties of the cue, which can be processed without extensive, and thus time-consuming, top–down control (14–16). The early onset of the cueing effect is also consistent with neurophysiological evidence demonstrating that neural signatures of covert attentional shifts can be detected in the thalamus and early visual cortices as early as ∼50 ms post-cue in non-human primates performing Posner-like paradigms (14, 17, 18). Converging evidence from human gaze-cueing studies further supports this interpretation in showing reliable cueing effects for non-predictive eye-gaze cues at short SOAs (see McKay et al. (19) and Chacón-Candia (20), for meta-analytic reviews). Although in most prior studies the shortest SOAs tested ranged between 100 and 150 ms (e.g. 21–27), at least one study explicitly testing a 50 ms SOA reported a robust gaze-cueing effect (28). Comparable early effects have also been documented in rhesus monkeys (Marciniak et al., 2015; Shafiei et al., in press). Perhaps the strongest evidence for a reflexive attentional mechanism comes from studies in which human participants were explicitly informed that the target was more likely to appear at the location opposite to the location cued by the gaze stimulus. Despite this counter predictive instruction, reliable gaze-cueing effects persisted at short SOAs (<200 ms), indicating that early gaze-following cannot be fully suppressed by top–down expectations (29, 30). A similar temporal pattern has been observed in rhesus monkeys: gaze cues exerted a robust influence on attention at short SOAs (50–200 ms), whereas at longer SOAs (≥300 ms) monkeys were able to suppress gaze-following and localize targets in accordance with learned task contingencies (31). Together, these findings support the view that eye-gaze cues engage an early, reflexive attentional orienting mechanism that operates prior to, and largely independently of, voluntary cognitive control.

A limitation of the present study is that we examined only human and not also monkey observers on their ability to follow the gaze of representatives of the respective other species. As we found that humans are much more proficient at eye gaze-following compared to rhesus monkeys, it may be unsurprising that they can flexibly generalize this ability to interpret gaze cues even when the eye morphology is unfamiliar or atypical, such as the rhesus monkeys’ eyes. Assuming that there is a direct link between sensitivity to eye information and the ability to cross the species boundary, one might speculate that rhesus monkeys may be less able to generalize across heterospecific eye configurations. Addressing this issue will require systematic behavioral evidence from NHPs tested with heterospecific eye-gaze cues, which remains sparse, even though a growing body of comparative morphological work suggests that—despite clear absolute differences in ocular morphology across species—key features relevant for gaze cue visibility (e.g., relative contrast) may not differ substantially. Moreover, it remains poorly understood that how facial context modulates gaze-following, as gaze perception likely reflects an integration of eye cues with surrounding facial information rather than the eyes alone. In other words, even identical eye morphologies may carry different behavioral relevance depending on the surrounding facial embedding and species-specific configural cues. An important next step therefore is to test how gaze-following changes when eye morphology and facial context are dissociated—for example, by embedding human eyes within a rhesus monkey face (and vice versa). For this purpose, computer-generated avatars —such as those used in the present study—will be critical, as they allow controlled manipulation of eye and face properties while preserving realism and minimizing low-level confounds.

A particularly intriguing question arising from the present findings is why humans are markedly more proficient than non-human primates (NHPs) at extracting spatial cueing information from very subtle eye-gaze signals. Given the broad similarities in the organization of the visual and oculomotor systems of diurnal old world primates including rhesus monkeys and humans (32– 34), it is unlikely that this difference can be attributed to fundamental neurobiological advantages in humans. Instead, learning through experience may account for humans’ higher sensitivity. Humans are exposed to substantially greater opportunities for eye-gaze interaction throughout development than most non-human primates. Arguably humans experience substantially greater opportunities for mutual eye-gaze contact across development than most non-human primates (35–37). From birth onward, human caregivers actively seek eye contact with infants to engage attention (38), and common caregiving practices—such as face-to-face holding (39), breastfeeding (40), and infant carrying—provide frequent opportunities for sustained mutual gaze. In contrast, in many non-human primate species, infants are typically carried on the mother’s back or ventrum during quadrupedal locomotion (41), a posture that likely limits opportunities for sustained face-to-face interaction and prolonged eye contact (42, 43). Later in development, humans routinely exchange eye contact in everyday social interactions as an affiliative signal, a cue for turn-taking, and an indicator of communicative intent, and brief eye contact is generally tolerated even in the absence of prior social bonding (44, 45). By contrast, in most non-human primate species, direct eye contact outside affiliative contexts such as grooming is often avoided, as it can be perceived as threatening and is frequently associated with aggression, dominance, or challenge displays, particularly when prolonged (2, 46, 47). Consequently, exposure to eye-gaze cues is likely far more constrained in NHPs than in humans throughout development and adulthood. Such differences in social experience may provide a developmental basis for the enhanced sensitivity and flexibility of human gaze processing, enabling humans to reliably extract spatial information from subtle eye-gaze cues. Nevertheless, we cannot exclude the possibility that humans’ greater perceptual sensitivity to eyes is already available from an early developmental stage (potentially driven by currently unknown differences in low-level visual processing), which in turn could strengthen the contribution of eye information to subsequent social-cognitive development.

In conclusion our results demonstrate a clear quantitative difference in gaze-following capabilities of humans and rhesus monkeys: humans can reliably extract gaze direction from small-amplitude cues that fail to drive gaze cuing in monkeys, despite both species showing qualitatively similar gaze-following capabilities under optimal conditions. Moreover, the near-equivalence of gaze-cuing effects elicited by human and monkey avatars in humans argues against the relevance of species-specific differences in eye morphology and coloration/contrast. Instead, the data suggest that the cross-species difference may reflect differences in perceptual sensitivity to subtle gaze signals rather than differences in the visual features of the eyes themselves.

## Materials and Methods

All procedures complied with the Declaration of Helsinki and were approved by the Ethics Committee of the University Hospital Tübingen. Participants provided written informed consent after receiving written and oral study information and received monetary compensation for their participation.

### Participants

A total of 28 neurotypical individuals were initially recruited through round-invitation emails sent to students and employees of the University of Tübingen. The following criteria were used for the initial recruitment of eligible participants: (1) availability to attend three experimental sessions, each lasting approximately 90 minutes; (2) no history of attention-deficit or attention-related disorders (e.g., ADHD); (3) no known color-vision deficiencies; (4) no sensitivity to flashing or flickering lights; and (5) absence of claustrophobia. During the first experimental session, participants’ ability to comply with the task requirements was evaluated which led to the exclusion of five participants (17.9%): three (10.7%) due to poor eye-tracking data quality caused by frequent blinking or difficulty maintaining their head stable, one (3.6%) due to inconsistent perception of luminance during the initial perceptual threshold estimation phase—likely related to uncorrected ametropia—and one due to an incomplete dataset following discontinuation of participation after the first session for personal reasons. None of the participants who completed all three experimental sessions were excluded from the final dataset. The final sample therefore comprised 23 participants (age 28.7 ± 9.81 years; 14 female). All participants reported normal or corrected-to-normal vision using prescribed glasses.

### Experimental procedure

The experiment consisted of three distinct phases: training, perceptual threshold estimation, and gaze-cueing paradigm. These were completed in three 90-min sessions, conducted on separate days, on average 3.23 days (SD = 4.05, range = 17 days) apart. The first session began with training, followed by the perceptual threshold estimation. The gaze-cuing task was mainly conducted in the 2^nd^ and 3^rd^ sessions. All experiments were conducted in a sound-attenuated booth which was dimly lit, with ambient lighting adjusted and controlled to maintain low illumination throughout the session. As shown in figure 3a, participants were seated in a height-adjustable chair with their head partially stabilized using an adjustable chin and forehead rest to enhance eye-tracking accuracy. Participants’ arms rested comfortably on a desk positioned in front of them. Visual stimuli were presented on a 24-inch LCD monitor (BenQ XL2411-B; 1920 × 1080 pixels, refresh rate 60 Hz) mounted at a viewing distance of 60 cm, with participants’ neutral line of sight aligned to the center of the display. A transparent plexiglass panel was positioned parallel to the screen at 14 cm in front of it (figure 3a). Two LEDs were affixed to the panel, positioned symmetrically at ±9.3° from the panel center along the horizontal meridian, aligned with the center of the screen. Eye movements were recorded using a desktop-mounted, video-based eye-tracking system (EyeLink® 1000 Plus, SR Research Ltd., Canada) with a sampling rate of 2 KHz. Stimulus presentation and eye-tracking data acquisition were controlled using the Neurological Recording and Experiment Control (NREC), an in-house developed software suite (for more details visit: https://nrec.neurologie.uni-tuebingen.de).

**Figure 3.**
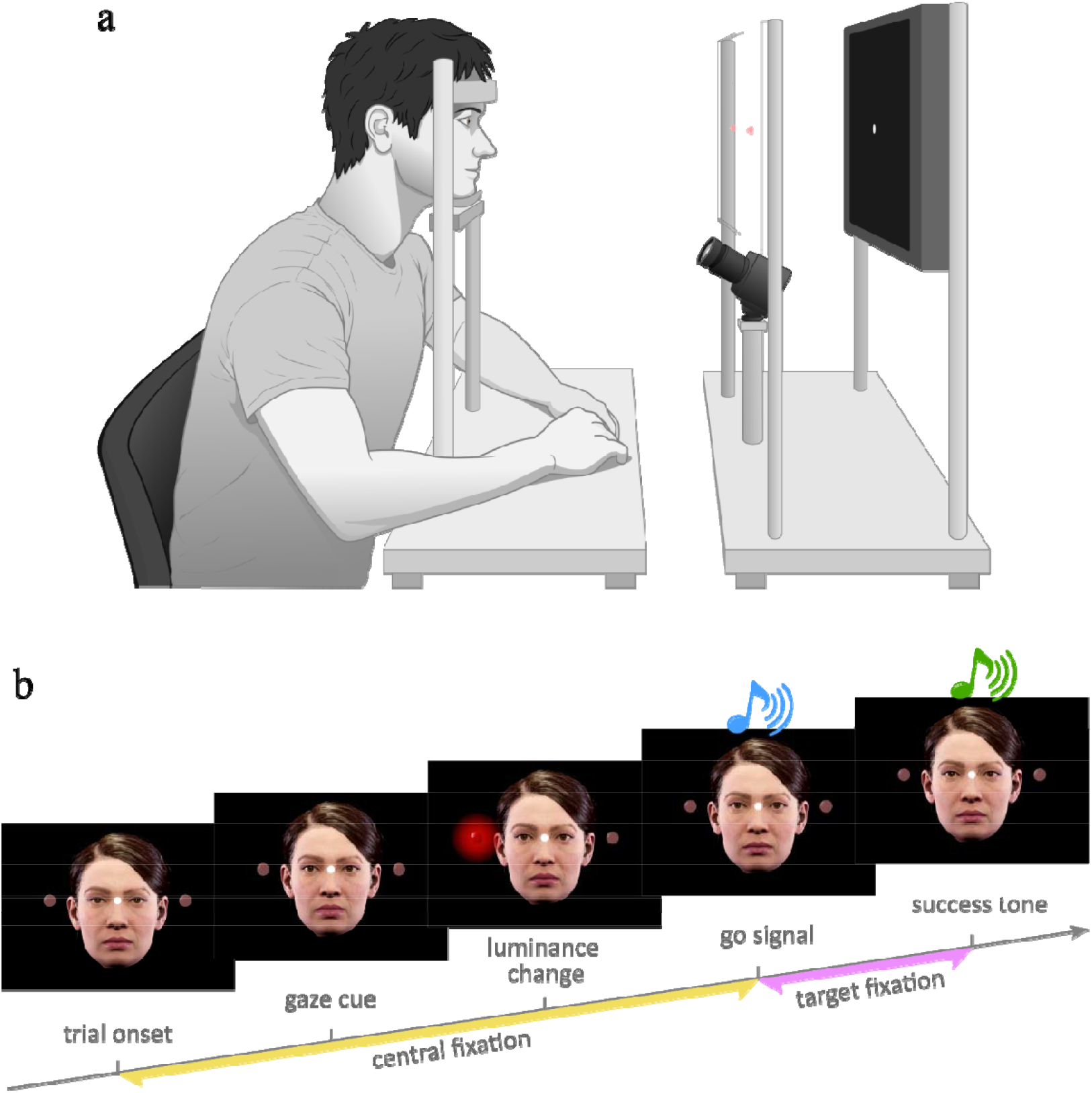
Experimental setup **(a)** and gaze-cuing paradigm **(b).** Faces shown in panels a and b are either illustrated (a) or generated entirely using a 3D graphics engine and are synthetic stimuli, not based on or derived from any real person.

### Phase I: training

The first experimental session began with a training phase consisting of 50 practice trials, preceded by a verbal explanation of the task. Prior to training, eye movements were calibrated using a standard nine-point calibration procedure, in which participants fixated nine points displayed sequentially across the screen in a 3×3 grid spanning 15° in width and 10° in height. This procedure allowed the eye tracker to accurately map measured eye positions to known screen locations. Each trial began with the presentation of a central fixation point (CF; white disk, 0.2° in diameter) on an otherwise black screen (figure S1a). Participants were instructed to maintain fixation on the CF to initiate the trial. Fixation was monitored using an invisible 5° × 5° window centered on the fixation point; leaving this window before the go signal resulted in immediate trial abortion. After a fixed interval of 500 ms, a salient light flash (duration: 300 ms) was presented by one of the two LEDs, thereby defining the target LED. Following a 500-ms delay after light offset, an auditory “go” signal was delivered. Participants were required to maintain fixation on the CF until the go signal and were instructed to report the target as quickly and accurately as possible by executing a saccade toward the corresponding LED upon hearing the signal. If the selected LED corresponded to the target, a distinct auditory feedback was provided indicating a correct response; incorrect responses were not followed by feedback. After an intertrial interval (ITI) of 700–1500 ms, the next trial began. During the training phase, verbal feedback was provided by the experimenters to facilitate understanding the task elements and performance. Once participants demonstrated consistent correct performance across several consecutive trials, verbal feedback was withheld until the end of training. If the learning criterion was not met after the initial 50 trials, training was extended by up to two additional blocks of 50 practice trials. Participants who failed to meet the criterion after a total of 150 training trials were excluded from further participation. Upon successful completion of training, participants proceeded to the subsequent phase, i.e. perceptual threshold estimation.

### Phase II: perceptual threshold estimation (PTE)

PTE was conducted to determine, for each participant, the LED luminance increase from baseline that enabled a 25% increase in detection accuracy relative to chance performance (50%), defined as the individualized point of subjective equality (PSE). The paradigm used in this phase closely matched the training paradigm, with luminance systematically varied on a trial-by-trial basis. Luminance levels ranged from near-chance detection to near-ceiling performance and were adaptively selected using a parameter estimation by sequential testing (PEST) procedure (48), which adjusted luminance intensity based on participants’ responses to converge on the target accuracy level of 75%. In brief, each trial began with the presentation of a central fixation point (CF) superimposed on a colored monkey-head silhouette against a black background (figure S1b). This background configuration was chosen to match overall screen luminance with that of the subsequent gaze-cueing phase, ensuring comparable light exposure across paradigm which critically impacts luminance perception. After a fixed delay of 675 ms, one of the two LEDs, selected at random, briefly increased in luminance for 300 ms. This delay corresponded to the midpoint of the range of delays (550–900 ms) used in the gaze-cueing task. The go signal followed 500 ms after LED offset, after which participants had up to 1500 ms to initiate an indicative saccadic response. Correct responses were confirmed by an auditory feedback tone followed by an ITI of 700–1500 ms.

PSE reliability was evaluated based on the shape of the psychometric function describing detection accuracy (with the lower asymptote fixed at chance level) to luminance intensity. This evaluation was performed immediately after completion of the perceptual threshold estimation (100 trials), while participants took a 5-minute break. An excessively steep transition between the lower and upper asymptotes was interpreted as insufficient resolution of luminance levels, whereas failure of the function to reach an upper asymptote indicated disproportionate task difficulty. To adjust LED visibility, we reduced LED luminance using neutral density (ND) filters placed over the LEDs and the ambient illumination within the testing booth. Pilot testing showed that reliable perceptual threshold estimation required attenuating the LEDs with one to three layers of ND filters to adequately modulate task difficulty. Once an appropriate filter was identified for an individual participant, it was kept constant across all subsequent experimental sessions. The resulting luminance value corresponding to 75% detection accuracy was subsequently used in the gaze-cueing phase of the experiment. If a reliable PSE could not be established after the initial 100 trials, LED visibility was adjusted and participants completed an additional 100 trials. This procedure was repeated for up to three rounds. Participants were excluded if a reliable PSE could not be obtained after the third round.

### Phase III: gaze-cuing paradigm

This paradigm builds on the spatial cueing task of Posner (49), in which a pre-cue shifts attention to a particular location. Here the other’s gaze served as pre-cue that guided attention to one of two LEDs, right and left of the fixation spot with one of the two exhibiting a luminance change to be detected by the experimental subject. The luminance could change at the LED cued by the demonstrator’s gaze or at the other LED. Gaze cues were provided by highly realistic pseudo-3D avatars of human and rhesus monkey heads respectively. The monkey head avatar has previously been shown to evoke behavioral responses in rhesus monkeys that do not differ from those elicited by videos of natural conspecifics (50). In addition, gaze cues offered by the same avatar have been shown to reliably elicit gaze-following behavior in monkeys (51). We used small-amplitude eye-gaze shifts identical to those described in Shafiei et al. (51). For the present study, gaze cues demonstrated by human head avatars were generated using Unreal Engine 5 (52) and CapCut (53). We carefully ensured that the gaze cues of the human avatars closely matched those of the monkey avatar. Specifically, saccade duration, speed, and amplitude, as well as eye size (comprising visible sclera and iris) and overall facial brightness, were matched across avatars. The accuracy of these adjustments was further verified using ImageJ (54), a free, open-source image analysis software widely used in science to view, measure, and process images. The avatars executed horizontal eye movements of 7.7° (when viewed from a distance of 60 cm), directed toward either the left or right LED (Figure 4). All participants were tested using gaze cues demonstrated by the monkey avatar. In addition, each participant was tested with one of two human avatar variants (male or female), chosen based on the participant’s subjective preference. Each avatar variant was tested in a separate block, with block order counterbalanced across participants. Each block consisted of 10 runs of 100 trials. Participants were allowed to take short breaks between runs.

**Figure 4.**
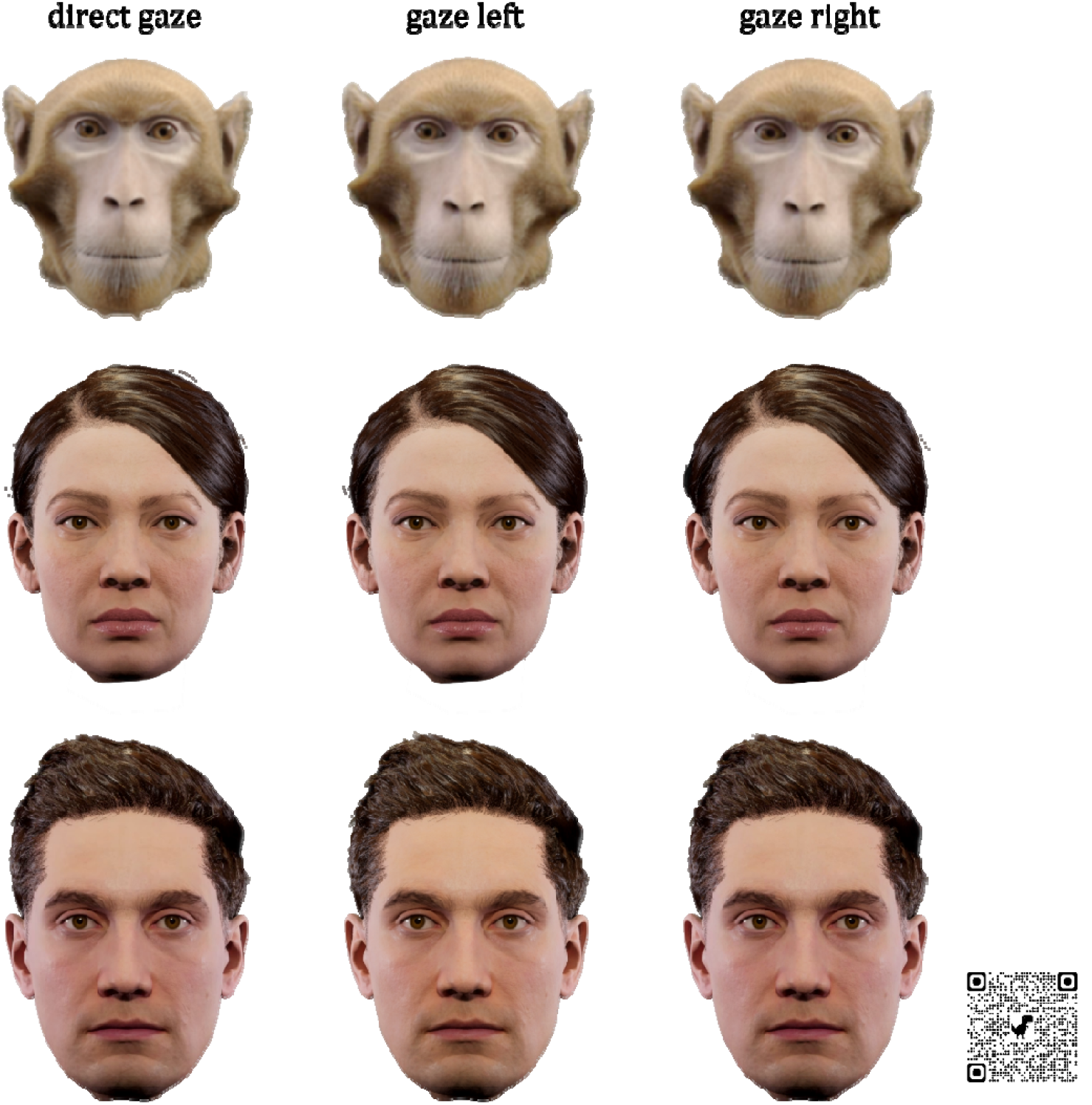
Snapshots of the avatars showing direct gaze (left) and gaze shifts to the left (middle) and right (right). Videos of the gaze cues are available at https://doi.org/10.6084/m9.figshare.31014625 or scanning the QR code. Human face stimuli were generated entirely using a 3D graphics engine and did not correspond to, or derive from, any real person.

Each trial in the main behavioral paradigm began with the presentation of a forward-facing avatar gazing straight with a central fixation point (CF) located on the bridge of its nose (Figure 3b). The CF was located midway on the horizontal axis connecting the two LEDs which exhibited the same level of baseline illumination. To initiate a trial, participants were required to maintain fixation within an invisible 5° × 5° window centered on the CF for 500 ms. Following successful fixation, the avatar shifted its gaze toward either the left or right LED, selected at random, and maintained this gaze direction until the end of the trial (figure 4). After a variable delay from the gaze shift onset, the luminance of one of the LEDs increased briefly for 300 ms, identifying the LED as the target in the given trial. The interval between the gaze shift onset and the luminance change is referred to as the stimulus onset asynchrony (SOA). The target LED (left or right) was selected at random (50% share each) and independently of the avatar’s gaze direction, ensuring that the gaze cue was uninformative with respect to target location. The LED luminance stepped from its baseline level to the individualized value determined during the perceptual threshold estimation phase that predicted 75% detection accuracy; this increase is referred to as ΔLuminance. Five hundred milliseconds after the offset of the ΔLuminance, the auditory go signal was presented. Participants were instructed to report the target LED by making a saccade toward it. Responses were required within 1500 ms following the go signal, after which the trial was aborted. Correct responses were indicated by an auditory feedback tone, whereas incorrect responses (saccades toward the non-target LED) were not. After either a correct or incorrect response, an ITI of 700– 1500 ms followed, during which the avatar’s gaze returned to the neutral position. Trials were self-paced, allowing participants to initiate the next trial by fixating on the CF. Before the experiment, participants received verbal instructions explaining the task that and learned that they should identify the LED whose luminance changed. They were also told that the gaze cue was irrelevant for identifying the target LED, i.e. that the gaze-cue did not reliably predict the target LED. Eye movements were then once again calibrated using the standard nine-point calibration procedure.

To distinguish between endogenous and exogenous sources of attention, we examined the temporal profile of the cueing effect by including five stimulus onset asynchronies (SOAs): 50, 100, 200, 300, and 400 ms. One SOA was randomly assigned to each trial, with equal representation of all SOAs within each run. In total, there were 20 trial types defined by the combination of gaze direction (left, right), target LED location (left, right), and SOA (five levels). Each trial type was repeated five times per run, with trials randomly interleaved. Because our primary interest was luminance detection accuracy in gaze-congruent (i.e., target LED located in the direction of the avatar’s gaze) versus gaze-incongruent trials, the four gaze-direction–by– LED-location combinations were collapsed into two congruency conditions. This resulted in 10 trials per congruency condition per SOA per run for each participant.

### Statistical Analysis

Data preprocessing was conducted using a custom-written script in MATLAB (R2023a)(55). Statistical data analysis and visualization were performed in R (version 4.4.2, 2024-10-31 ucrt)(56). The detection of response saccades and computation of their metrics were made using SAL (Saccade Analysis Lab), a custom-made MATLAB-based algorithm (Version: 9.10.0.1710957 (R2021a) Update 4)(57) adapted from the method described by Nyström and Holmqvist (58). The full SAL codebase is openly available for reproducibility at: https://github.com/shafieimasih/SAL.

Estimation of PSE was performed by modelling hit rate as a function of ΔLuminance level using a logistic psychometric function using *psignifit* package in MATLAB (59), with the lower asymptote fixed at 50% to reflect chance-level performance.

Behavioral performance was analyzed at the trial level using generalized linear mixed-effects models (GLMMs) implemented in R (lme4 package (60)). The dependent variable was hit probability (correct vs. incorrect response), modeled with a binomial error distribution and a logit link function. For the primary analysis, fixed effects included avatar type (human, monkey), congruency (congruent, incongruent), stimulus onset asynchrony (SOA; 50, 100, 200, 300, 400 ms), and Congruency-by-SOA and avatar-type-by-congruency interactions. Random effects were specified to account for repeated measurements within participants, including random intercepts for subjects and random slopes for congruency by subject. This random-effects structure allowed baseline performance and the magnitude and direction of the congruency effect to vary across individuals. Model convergence and fit were verified by inspection of residuals and random-effects estimates. Statistical significance of fixed effects was further assessed using *Wald z* tests. Effect sizes are reported as regression coefficients (*β*), standard errors (*SE*), *z* values, and associated *p* values. To facilitate interpretation of the congruency effect, odds ratios (ORs) were computed from model estimates. Post-hoc comparisons were conducted to examine the congruency effect separately for each avatar type and SOA. Estimated marginal means and contrasts were computed deploying the *emmeans* package (61), with *p* values adjusted for multiple comparisons using the Benjamini-Hochberg method.

To assess whether participant sex, avatar presentation order, or human avatar sex influenced task performance or the gaze-cuing effect, we conducted a series of additional generalized linear GLMMs with a binomial error distribution and logit link function. In these models, hit probability (hit vs. miss) was modeled as a function of avatar type (human vs. monkey), SOA, gaze congruency, and their interactions, while including each control variable of interest as an additional fixed effect. Participant sex (male vs. female) and avatar presentation order (monkey-first vs. human-first) were included as between-subject factors, whereas human avatar sex (male vs. female avatar) was included as a within-subject factor in analyses restricted to human avatar type. All models included random intercepts and random slopes for congruency by subject to account for individual differences in baseline performance and gaze-cuing sensitivity. Models were fitted using maximum likelihood estimation with Laplace approximation, and significance of fixed effects was assessed using Wald z tests.

For cross-species comparisons, odds ratios derived from the human data were compared with corresponding ORs from our previous rhesus monkey study (51), which employed the identical experimental paradigm and the same small-amplitude monkey-avatar gaze cue. ORs provide a scale-free measure of effect size that enables quantitative comparison of the gaze-cuing effect across studies and species. All statistical tests were two-tailed, and the significance threshold was set at α = .05.

## Acknowledgments

We thank Dr. Friedemann Bunjes and Peter W. Dicke for their invaluable technical assistance.

